# Sequence-to-structure dependence of isolated IgG Fc complex biantennary N-glycans: A molecular dynamics study

**DOI:** 10.1101/392001

**Authors:** Aoife M Harbison, Lorna P Brosnan, Keith Fenlon, Elisa Fadda

## Abstract

Fc glycosylation of human immunoglobulins G (IgGs) is essential for their structural integrity and activity. Interestingly, the specific nature of the Fc glycoforms is known to modulate the IgG effector function. Indeed, while core-fucosylation of IgG Fc-glycans greatly affects the antibody-dependent cell-mediated cytotoxicity (ADCC) function, with obvious repercussions in the design of therapeutic antibodies, sialylation can reverse the antibody inflammatory response, and galactosylation levels have been linked to aging, to the onset of inflammation, and to the predisposition to rheumatoid arthritis. Within the framework of a structure-to-function relationship, we have studied the role of the N-glycan sequence on its intrinsic conformational propensity. Here we report the results of a systematic study, based on extensive molecular dynamics (MD) simulations in excess of 62 µs of cumulative simulation time, on the effect of sequence on the structure and dynamics of increasingly larger, complex biantennary N-glycoforms, i.e. from chitobiose to the larger N-glycan species commonly found in the Fc region of human IgGs. Our results show that while core fucosylation and sialylation do not affect the intrinsic dynamics of the isolated (unbound) N-glycans, galactosylation of the α(1-6) arm shifts dramatically its conformational equilibrium from an outstretched to a folded conformation. These findings are in agreement with and can help rationalize recent experimental evidence showing a differential recognition of positional isomers in glycan array data and also the preference of sialyltransferase for the more reachable, outstretched α(1-3) arm in both isolated and Fc-bound N-glycans.

## Introduction

N-glycosylation of the immunoglobulin G (IgG) fragment crystallizable (Fc) region is essential for its structural stability and function^1-4^. The sequence and branching of the Fc N-glycoforms, bound at the highly conserved Asn 297 in both CH_2_ domains of the Fc region, strongly affect the antibody-mediated effector function^5-7^, by modulating the binding affinity of Fcγ receptors (FcγRs), thus the antibody-mediated immune response ^8-9^. In this context the effects of core-fucosylation, sialylation and of galactosylation are particularly interesting. Between 81% and 98.7% of the Fc N-glycans in human IgGs are core-fucosylated^10^. Even though it may appear as a subtle change to the glycan sequence, core fucosylation, where fucose is α(1-6) linked to the chitobiose core of the complex N-glycan, greatly affects the IgGs antibody-dependent cell-mediated cytotoxicity (ADCC) function. More specifically, a strongly enhanced ADCC corresponds to non-fucolylated F_c_ N-glycan species^6, 11-16^. This information has found wide interest and applications in cancer immunotherapy, especially in regards to engineering non-fucosylated antibodies with higher efficacy^7, 13, 17-18^. The molecular basis for this phenotype is not entirely clear. It has been linked to the stronger binding^7, 19-20^ between IgG and the FcγRIIIa, due to a more effective contact between the IgG and FcγRIIIa glycans in the absence of core fucose^19^. No significant structural changes in the IgG structure have been detected in function of the presence or absence of fucose^11^. Sialylation of the Fc glycans is known to reverse the antibody pro-inflammatory effect into anti-inflammatory^21^. Only about 20% of core-fucosylated biantennary Fc N-glycans are sialylated^10^, meanwhile the majority of Fc N-glycans in human IgGs are galactosylated, with neutral glycans without galactose slightly below 40%, neutral glycans with one terminal galactose slightly above 40%, and neutral glycans with two terminal galactoses contributing to 20% of the neutral IgG glycome^10^. The abundance of galactosylated glycoforms has been directly linked to aging, with decreasing levels correlated with aging, and to immune activation^10, 22-24^. Notably, the risk of developing rheumatoid arthritis is correlated with low levels of galactosylation^25-27^.

The phenotypes linked to different Fc N-glycan sequences are likely to be determined by the modulation of the interaction of the IgG with cell surface receptors and/or lectins, which is a difficult topic to address as a whole due to the complexity of the systems involved. NMR spin relaxation data provide evidence that despite the close contact with the protein, both arms of the N-glycans at the IgG Fc remain flexible and accessible ^28^, suggesting that the intrinsic conformational propensity of the Fc N-glycan in function of its sequence may play a role in regulating the molecular interaction with the cell surface receptors. Therefore, to understand the implications on the N-glycans structural and dynamics of the size and sequence, we conducted a complete conformational study by extensive molecular dynamics (MD) simulations of progressively long complex biantennary Fc N-glycans most commonly expressed in human IgGs ^10, 29^. All the glycoforms we have analysed in this work are shown in **Figure 1**. Because of the complexity of the N-glycans dynamics and the high flexibility of α(1/2-6) linkages that can access many different conformational states, we chose a sampling method based on single (conventional) MD trajectories, ran in parallel, all started from different combinations of α(1/2-6) conformers, namely 3 trajectories for one α(1-6) linkage, 9 for two α(1-6) linkages, 12 for one α(1-6) and one α(2-6) linkages, 24 for two α(1-6) and one α(2-6) linkages, and finally 72 for two α(1-6) and two α(2-6) linkages, for a total cumulative sampling time in excess of 62 µs. This approach allowed us to systematically and directly sample by construction regions of the potential energy surface corresponding to rare conformations of the different α(1/2-6) linkages and to assess their relative stability. Our results show that a) the highest conformational flexibility concern primarily the α(1-6) arm, while the α(1-3) arm remains relatively flexible but mostly extended, b) core fucosylation and sialylation do not affect the conformational equilibrium of the α(1-6) arm in the isolated glycan in solution, meanwhile c) galactosylation of the α(1-6) arm alone greatly shifts the conformational propensity of the arm from outstretched, to folded over the chitobiose core. These findings provide important insight into the differences in the molecular recognition of biantennary complex N-glycans by enzymes and lectins in function of their sequence.

**Figure 1.**
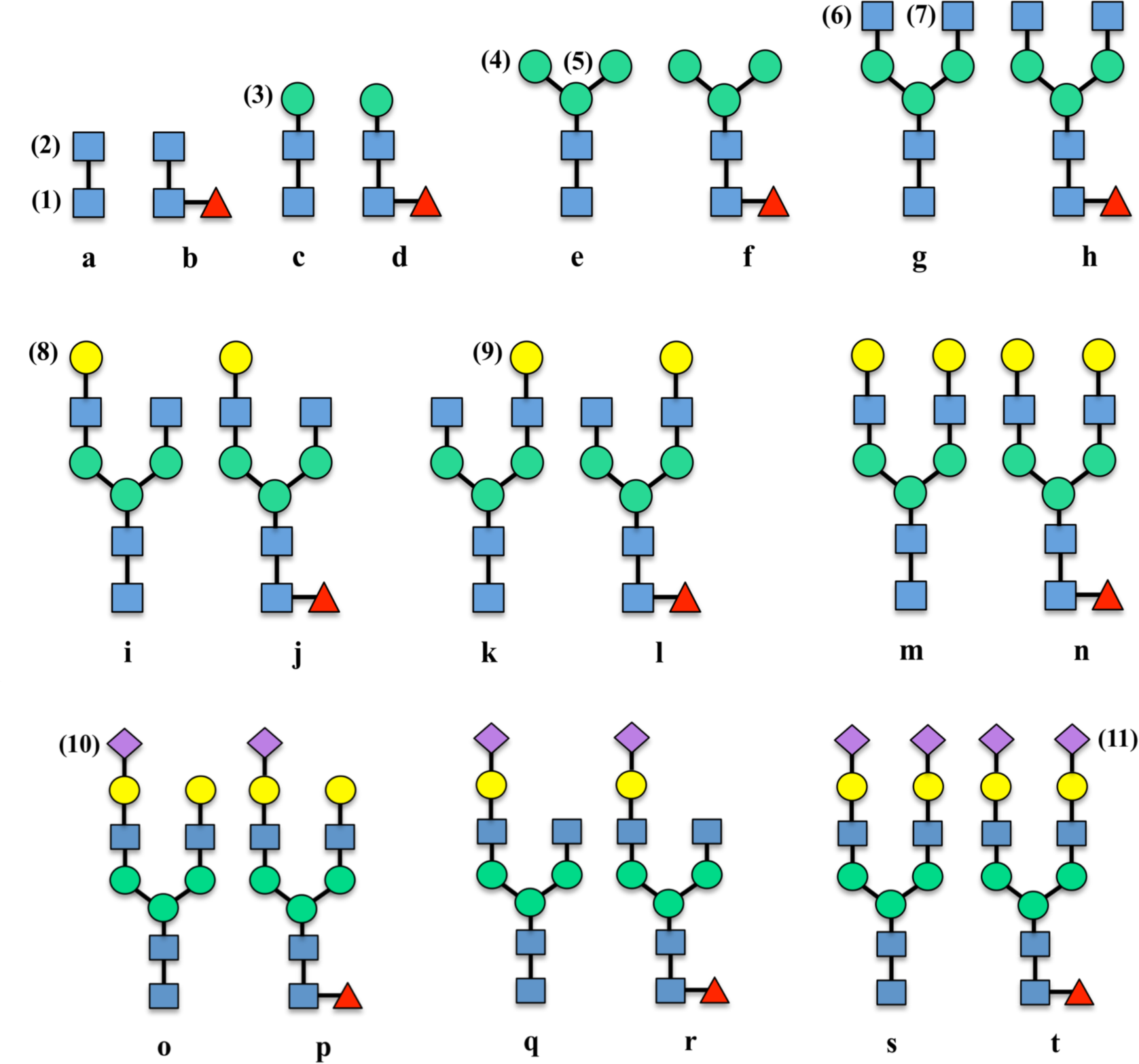
Schematic representations of all N-glycans, fucosylated and non-fucosylated in pairs, analysed in this study. Letters are used as shorthand notation for the identification of each sugar. Numbers are used to identify residues. The graphical representation follows the guidelines indicated in ^30^.

## Computational Methods

All glycans were built with the carbohydrate builder tool on GLYCAM-WEB (http://www.glycam.org). All combinations of rotamers for the α(1/2-6) linkages have been considered as starting structures for the MD simulations, namely 3 trajectories for one α(1-6) linkage, 9 for two α(1-6) linkages, 12 for one α(1-6) and one α(2-6) linkages, 24 for two α(1-6) and one α(2-6) linkages, and finally 72 for two α(1-6) and two α(2-6) linkages. The GLYCAM-06h-12SB version of the GLYCAM06 force field^31^ was used to represent the carbohydrate atoms, TIP3P parameters ^32^ were used to represent water, while amber99SB parameters^33^ were used for the counterions, added in a number sufficient to neutralize the charge in the case of sialylated species.

All simulations were run with versions 12 and 16 of the AMBER molecular simulation package^34^. Dispersion interactions were cutoff at a distance of 13 Å. Electrostatics interactions were treated with Particle Mesh Ewald (PME). A constant pressure of 1 atm was maintained by isotropic position scaling with a relaxation time of 2 ps, while a constant temperature of 300 K was regulated by Langevin dynamics using a collision frequency of 1.0 ps^−1^. The SHAKE algorithm was used to restrain bonds with hydrogen atoms and an integration time step of 2 fs was used throughout. For each sugar, each starting structure was analysed for at least 250 ns, with exceptions of sugar D, which was analysed for 3 µs, because of its complex dynamics that will be discussed in the sections below, and sugar E for 500 ns on single trajectories. Further details on the minimization, equilibration and production phases are included as Supplementary Material. High performance computing (HPC) resources were provided by the Irish Centre for High-End Computing (ICHEC).

As an interesting note, because of the better scaling on our machines of v. 4.6.3 and 5.0.x of GROMACS (GMX) for the calculations on these relatively small systems, we ran some tests on the medium-sized sugar H, see **Figure 1,** and compared the results with the GLYCAM/AMBER set-up. The starting structure and parameter files obtained from the carbohydrate builder on GLYCAM-WEB were converted to GMX format with the AnteChamber PYthon Parser interfacE (*acpype.py*) tool^35^. It is important to note that in all GMX simulations the 1-4 scaling was re-set to “1” as required by the GLYCAM force field^31, 36^. Equal amount of sampling was done with both GMX and AMBER, preceded by a very similar set-up and equilibration protocols. Details of the GMX protocol are provided as Supplementary Material. The results indicate large differences in a(1-6) torsions populations between GMX and GLYCAM/AMBER, shown in **Tables S.1** and **S.2.** The reason for this may be problems in the transfer of torsional parameters from a GLYCAM/AMBER format to the GMX format. More specifically we found that simulations of sugar H with GMX do not reproduce the correct conformer populations, or give energetically unfavoured conformers, such as the <u>*tg*</u> in the core fucose α(1-6) linkage as the highest populated for sugar H2, see **Table S.2**.

## Results

The results below are presented in function of the different N-glycan linkages for clarity. Notable effects on the conformational propensity of different linkages determined by the N-glycan size and sequence are indicated within. All torsion angles discussed throughout correspond to the following nomenclature, ϕ (O_5_C_1_O_x_C_x_), ψ (C_1_O_x_C_x_C_x+1_), and ω (O_1_C_x_C_x_C_x+1_)^37^. The method we have chosen to number the monosaccharides and to name the different N-glycans is indicated in **Figure 1**. A summary of the results obtained as averages over all the simulations for all fucosylated and non-fucosylated species is shown in **Tables 1** and **2**, respectively. Results obtained for each system studied are shown in **Tables S.3-17**.

**Table 1.**
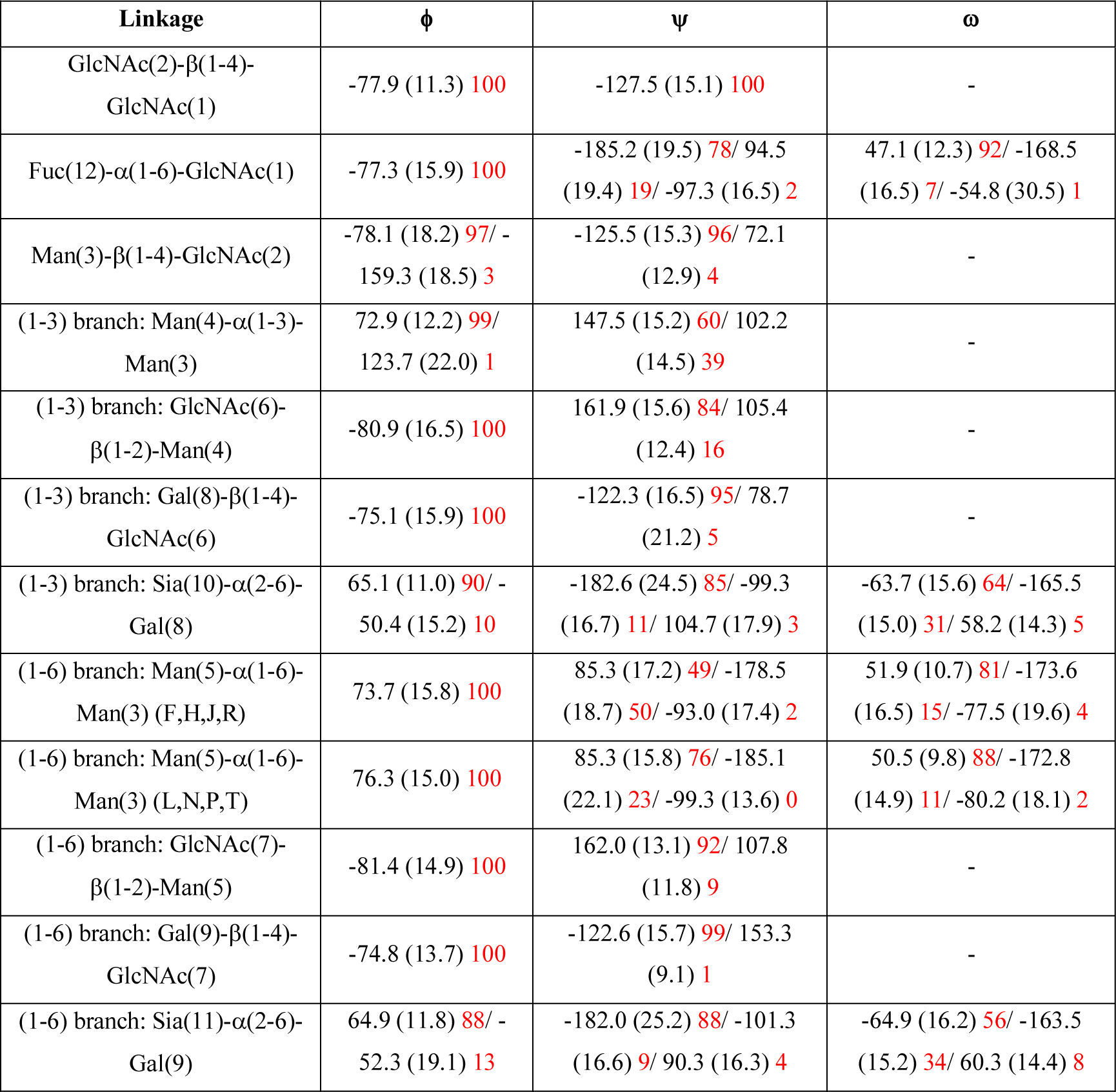
Conformational propensities of different linkages calculated for all <u>core-fucosvlated species</u> shown in **Figure 1**. The torsion angle values are shown in degrees and calculates as averages over all N-glycans. Data were collected and analysed at 100 ps intervals. Errors are shown in parenthesis and are averages of standard deviations calculated for each N-glycan. Relative populations (%) are indicated in red. All torsion angles discussed throughout correspond to the following nomenclature, ϕ (O5C1OxCx), ψ (C1OxCxCx+1), and ω (O6C6C5C4).

### GlcNAc(2)-β(1-4)-GlcNAc(1) linkage

As shown in see **Tables 1** and **2**, this linkage is conformationally rigid with only one rotamer populated in all the N-glycans analysed. The average ϕ angle values are −78.7° and −77.9°, while the ψ angle values are −130.8° and −127.5° for non-fucosylated and for fucosylated species, respectively. In the core-fucosylated species this conformation of the chitobiose favours the formation of a hydrogen bond between the O2 of the fucose and the NH of GlcNAc(2). No significant deviations from these torsion angle values have been observed, except in the case of the core-fucosylated tetrasaccharide sugar D, shown in **Figure 1**, where the GlcNAc(1) has been observed to change conformation from the more stable ^4^C_1_ to a ^1^C_4_ chair, causing a transition to a ψ angle of −74.0° for 13% of the time calculated over a 3 µs trajectory. Because this change in the ψ angle has been seen exclusively for sugar D and it is due to the transition to a different sugar conformation which reorients the GlcNAc(1) C5-C6 from an equatorial to an axial position, sugar D has not been included in averaging the values shown in **Table 1.** The conformational propensity of sugar D is discussed in detail in a dedicated subsection below.

### Fuc(12)-α(1-6)-GlcNAc(1) linkage

The flexibility of the α(1-6) linkage between the fucose and the GlcNAc(1) is unsurprisingly higher relative to the other core monosaccharides. Notably, the conformational space sampled is independent of the length or the size of the N-glycan. The highest populated α(1-6) conformer, i.e. ϕ −77.3° (100%), ψ −185.2° (78%), and ω 47.1° (89%), with relative average populations indicated in brackets, corresponds to a structure where the fucose O2 forms a hydrogen bond with the NH of GlcNAc(2). An example of this conformation is shown in the case of the dodecasaccharide sugar T in **Figure 2**. As the size of the glycan increases the fucose is also able to extend the hydrogen bond network by forming interactions with the GlcNAc(7) O3 when the α(1-6) arm is folded over the chitobiose core, as shown in **Figure 2**. A deviation from this conformation with 19% of the total population, consistently for all N-glycans, has the same ϕ and ω torsions, but a ψ value of 94.5°. In this conformer the fucose is not hydrogen bonded. A unique case where the ω torsion deviates from its <u>*gg*</u> stable configuration, i.e. ω of 47.1°, is found only in the case of the tetrasaccharide sugar D, where the reducing GlcNAc(1) ^1^C_4_ chair conformation has the C5 in an axial positions and the ω torsion to adopt a −64.5° value. This ω value corresponds to an energetically prohibited <u>*tg*</u> conformer when the GlcNAc(1) is in the standard ^4^C_1_ chair. The <u>*gt*</u> ω conformation, corresponding to an average value of −168.7°, has relative populations ranging between 3% (sugar T) and 15% (sugar F) in a pattern that doesn’t seem to be dependent on the branching, the length, nor on the sequence.

**Figure 2.**
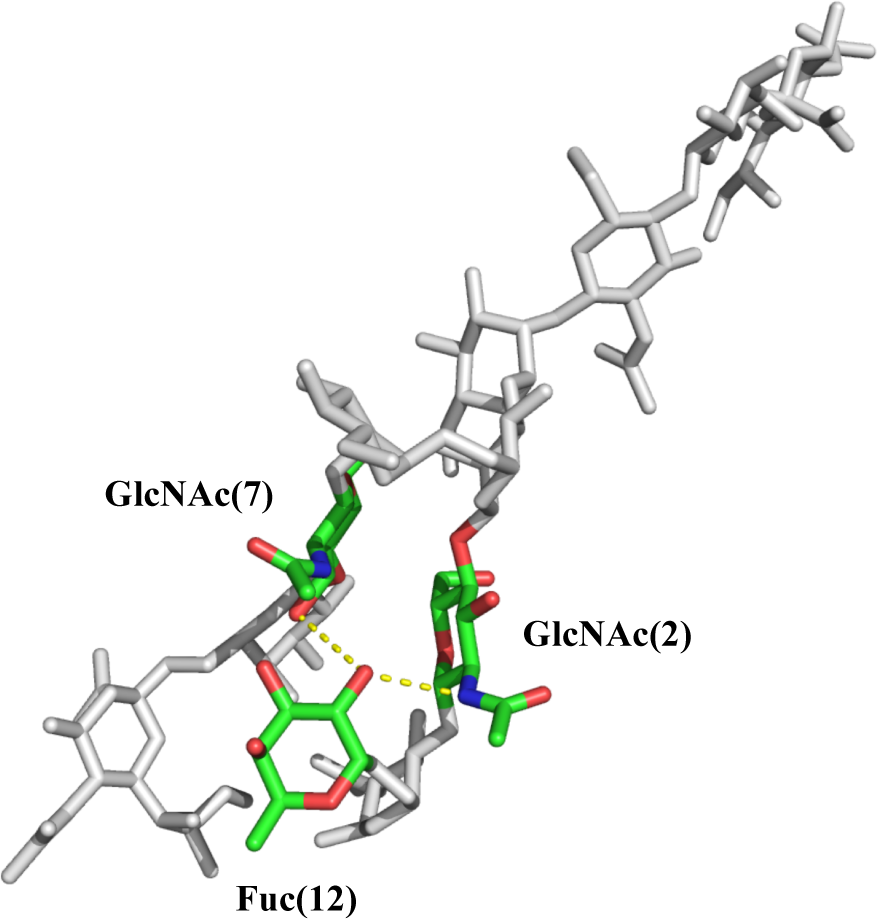
Hydrogen bond between the Fuc(12) O2 and the amide N of GlcNAc(2) and GlcNAc(7) in the dodecasaccharide sugar T is highlighted with yellow dots. Aside from Fuc(12) and GlcNAc(2) all other monosaccharides are shown in light grey. Note: the conformation shown, the highest populated in sugar T, has the α(1-6) arm folded over the chitobiose core. The image was generated with pymol.

### Man(3)-β(1-4)-GlcNAc(2) linkage

This linkage has one largely preferred conformation for both, fucosylated and non-fucosylated species, irrespectively of the length and sequence of the N-glycan. For non-fucosylated species ϕ and ψ have average values of −79.6° (97%) and −125.4° (96%), respectively. The relative populations calculated over the entire simulation time are indicated in parentheses. For core-fucosylated species ϕ and ψ have average values of −78.1° (97%) and −125.5° (96%), respectively. Conformational changes from this prevalent configuration amount to 3% to 4% of the whole cumulative simulation time and represent changes of ϕ to values around −160° and of ψ to values around 60° for both, non- and core-fucosylated species, see **Tables 1** and **2**. These ϕ and ψ excursions are not correlated to each other and confer a slight flexibility in terms of rotations of the plane containing the α(1-3) and α(1-6) branches around the chitobiose core. In the most stable conformation the planes containing the chitobiose and the trimannose core are parallel to each other.

### α(1-3) branch: Man(4)-α(1-3)-Man(3) linkage

The conformational dynamics of the α(1-3) branch at the trimannose level is not particularly complex and it is not affected by the core fucosylation. The ϕ torsion is prevalently found in a single conformation, 72.8° in 100% of all non-fucosylated species analysed and 72.9° in 99% of all core-fucosylated species analysed. The slight difference is due to the more complex dynamics of sugar P and sugar T, which both have sialylated α(1-3) arms, and for which in 5% and 1% of the sampling time the ϕ torsion value is an average of 123.7°, see **Table S.8**. The ψ torsion defines two distinct basins, identical in case of both fucosylated and non-fucosylated species. In case of non-fucosylated N-glycans the highest populated ψ value is 146.5° (61%) and the lowest is 102.3° (39%). In the case of fucosylated species the ψ values are, 147.5° with a relative population of 60% and 102.2 with a population of 39%. The remaining 1% is contributed by the more complex dynamics of sugar P, where the third ψ value corresponds to 182.0°. The two distinct conformations of this linkage confer a degree of flexibility to the α(1-3) branch with a range of movement shown in **Figure 3**.

**Figure 3.**
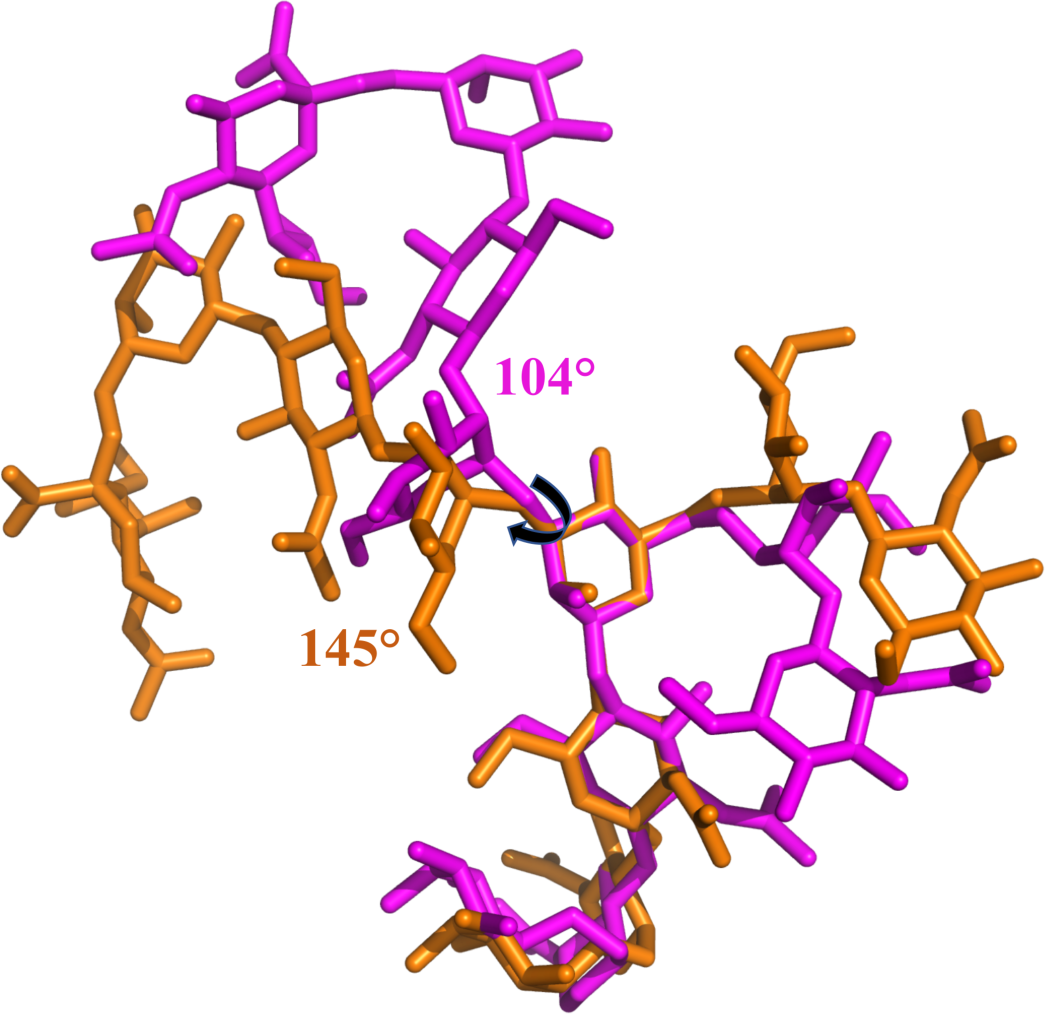
The conformational dynamics of the α(1-3) arm, shown here for sugar R as an example, is restricted to two basins corresponding to ψ values of 145° (orange) and 104° (purple). Rendering done with pymol.

### α(1-3) branch: GlcNAc(6)-β(1-2)-Man(4) linkage

The conformation of this linkage is restricted to two conformations, identical for both fucosylated and non-fucosylated species. In case of the non-fucosylated glycans the average ϕ value is −81.0°, while the ψ values are 161.8° (84%) and 105.4° (16%). In case of the fucosylated glycans the average ϕ value is −80.9°, while the ψ values are 161.9° (84%) and 105.4° (16%).

### α(8)-β-GlcNAc(6) linkage

This linkage is the least flexible in the branch with one conformation accounting for over 95% of the population of both fucosylated and non-fucosylated N-glycans. In case of the non-fucosylated glycans the average ϕ value is −75.7°, while the ψ value is −122.1° (95%). In case of the fucosylated glycans the average ϕ value is −75.1°, while the ψ value is −122.3° (95%). The 5% difference corresponds to excursions to ψ values around 75° for both fucosylated and non-fucosylated species, see **Tables 1** and **2.**

### α(1-3) branch: Sia(10)-α(2-6)-Gal(8) linkage

The linkage to the terminal sialic acid is the most flexible of the α(1-3) arm and all the conformations visited are independent of core-fucosylation, both in terms of torsion angle values and populations, see **Tables 1** and **2**. The most flexible torsion is the ω angle with values around −60° (65%) and −165° (30%). A small contribution of around 5% is given by a value of 60°.

### α(1-6) branch: Man(5)-α(1-6)-Man(3) linkage

The conformational dynamics of the Man(5)-α(1-6)-Man(3) linkage is the most interesting and the only one that has a clear dependence on sequence among the IgG N-glycans analysed here. Unlike the α(1-3) arm that has a relatively restricted dynamics, the α(1-6) arm can adopt two distinct conformations, one extended, or ‘outstretched’, with ϕ values around −180°, and one where the arm is folded over the chitobiose core, which we’ll refer to as ‘folded-over’, with ϕ values around 80°, see **Tables 1, 2** and **Figure 4.** In sugar E (F), with an α(1-6) arm terminating with Man, over 80% of the conformer have the arm in an outstretched conformation, see **Tables S.12** and **S.13.** The addition of the GlcNAc shifts this equilibrium, so for sugars G (H), I (J), and Q (R), where the fucosylated species are indicated in parenthesis, the outstretched and the folded-over conformations are equally populated, namely 45% (49%) for the folded-over conformation, and 55% (51%) for the outstretched conformation. Interestingly, galactosylation of the α(1-6) arm shifts this equilibrium further with a high majority of the galactosylated sugars, namely K (L), M (N), O (P) and S (T), in a folded-over conformation. The relative average populations are 74% and 76% in a folded-over conformation for non-fucosylated and fucosylated sugars, respectively, while 26% and 23% in an outstretched conformation for non-fucosylated and fucosylated sugars, respectively. This equilibrium is not affected either by the type of glycosylation in the α(1-3) arm, nor by the sialylation of the α(1-6) arm, and it does not depend on core-fucosylation either, despite the fucose in its most stable conformation interacts effectively through hydrogen bonding with the α(1-6) arm when folded-over.

**Table 2.** Conformational propensities of different linkages calculated for all <u>non-fucosvlated species</u> shown in **Figure 1**. The torsion angle values are shown in degrees and calculates as averages over all N-glycans. Data were collected and analysed at 100 ps intervals. Errors are shown in parenthesis and are averages of standard deviations calculated for each N-glycan. Relative populations (%) are indicated in red. All torsion angles discussed throughout correspond to the following nomenclature, ϕ (O5C1OxCx), ψ (C1OxCxCx+1), and ω (O6C6C5C4).

| Linkage | $\phi$ | $\psi$ | $\omega$ |
| --- | --- | --- | --- |
| GlcNAc(2)-β(1-4)-<br>GlcNAc(1) | -78.7 (11.1) **100** | -130.8 (15.7) **99**/ 69.0<br>(12.4) **3** | - |
| Man(3)-β(1-4)-GlcNAc(2) | -79.6 (20.6) **97**/ -157.9<br>(18.2) **1**/ 56.7 (9.1) **1** | -125.4 (15.4) **96**/ 66.8<br>(11.8) **3** | - |
| (1-3) branch: Man(4)-α(1-3)-Man(3) | 72.8 (12.1) **100** | 146.5 (13.9) **61**/ 102.3<br>(14.2) **39** | - |
| (1-3) branch: GlcNAc(6)-<br>$\beta$(1-2)-Man(4) | -81.0 (16.3) **100** | 161.8 (15.9) **84**/ 105.4<br>(12.3) **16** | - |
| (1-3) branch: Gal(8)-β(1-4)-<br>GlcNAc(6) | -75.7 (17.3) **100** | -122.1 (16.4) **95**/ 74.5<br>(17.6) **5** | - |
| (1-3) branch: Sia(10)-α(2-6)-Gal(8) | 65.1 (11.2) **89**/ -50.0<br>(14.7) **11** | -177.2 (22.9) **80**/ -100.3<br>(15.9) **18**/ 105.9 (15.8) **3** | -63.2 (15.5) **66**/ -164.3<br>(14.8) **31**/ 60.3 (13.1) **4** |
| (1-6) branch: Man(5)-α(1-6)-Man(3) (E,G,I,Q) | 73.2 (16.5) **100** | 82.1 (18.4) **45**/ -177.7 (18.6) **54**/ -95.6 (17.7) **2** | 51.8 (11.0) **80**/ -172.3 (17.0) **15**/ -75.7 (17.9) **5** |
| (1-6) branch: Man(5)-α(1-6)-Man(3) (K,M,O,S) | 75.1 (14.9) **100** | 82.9 (17.3) **74**/ -182.2 (21.9) **26**/ -101.8 (13.4) **1** | 58.4 (10.3) **99**/ -173.5 (18.5) **1** |
| (1-6) branch: GlcNAc(7)-β(1-2)-Man(5) | -79.8 (19.6) **100** | 162.1 (13.6) **90**/ 106.5 (12.4) **10** | - |
| (1-6) branch: Gal(9)-β(1-4)-GlcNAc(7) | -76.2 (15.1) **100** | -125.7 (15.4) **97**/ 120.3 (7.2) **3** | - |
| (1-6) branch: Sia(11)-α(2-6)-Gal(9) | 64.5 (11.2) **90**/ -50.7 (13.7) **10** | -181.0 (25.2) **88**/ -98.6 (17.9) **9**/ 106.5 (16.0) **3** | -62.6 (15.6) **60**/ -166.6 (14.3) **33**/ 58.8 (13.6) **7** |

**Figure 4.**
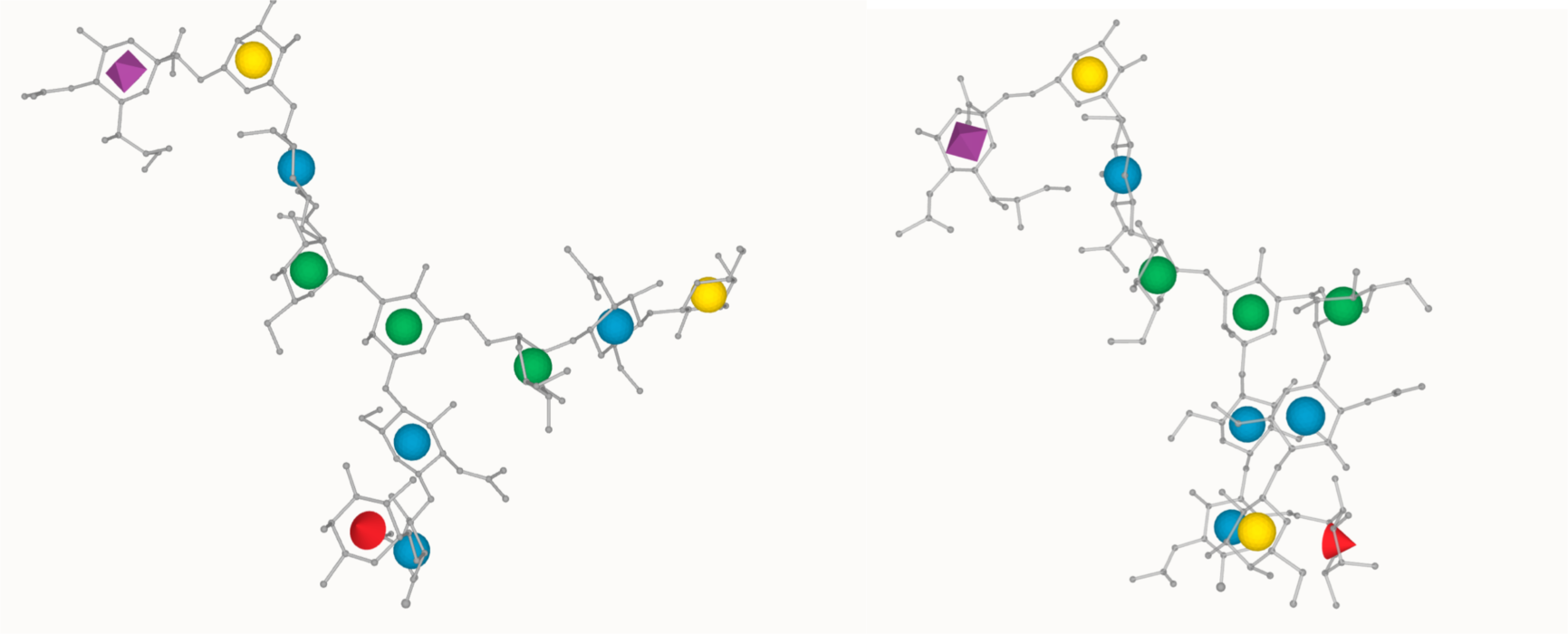
Two main conformations accessible to the α(1-6) arm represented for sugar P, chosen as an example. The ‘outstretched’ conformation is shown on the left and the ‘folded-over’ on the right. Rendering with LiteMol 3D SNFG^38^.

### α(1-6) branch: GlcNAc(7)-β(1-2)-Man(5) linkage

As seen in the case of the a(1-3) arm, the conformational dynamics of this linkage is largely restricted to ϕ and ψ values of −80° and 162°, respectively, both in case of fucosylated and non-fucosylated species, **see Tables 1** and **2**. The only significant alternative conformer accounting for 10% total contribution to the population is characterized by ψ values of 106° for non-fucosylated species and 108° for fucosylated species. The same linkage on the α(1-3) arm shows a similar conformational equilibrium, but with a slightly different 85:15 ratio.

### α(1-6) branch: Gal(9)-β(1-4)-GlcNAc(7) linkage

Similarly to the same linkage on the α(1-3) arm, the Gal(9)-β(1-4)-GlcNAc(7) torsional space is highly restricted to a single conformation with ϕ and ψ values around −75° and −125°, respectively, both in case of fucosylated and non-fucosylated species.

### α(1-6) branch: Sia(11)-α(2-6)-Gal(9) linkage

The conformational dynamics of the terminal sialic acid on the α(1-6) arm is virtually identical to the one observed for the terminal sialic acid on the α(1-3) arm. The linkage is highly flexible with, the ϕ angle predominantly around 65° (90%), with a ~ 10% contribution of −52°. The ψ angle has also a preferred conformation around −180° (88%). Small contributions of ψ (3-4%) have been also detected, both in case of fucosylated and of non-fucosylated species. The ω angle is quite flexible, with values of −65° as the preferred conformation (60%) and −162° (33%). A small contribution of 7% corresponds to an ω value of 60°, both in case of fucosylated and of non-fucosylated species.

### Sugar D: Man-β(1-4)-GlcNAc-β(1-4)-[α(1-6)-Fuc]-GlcNAc

The conformational dynamics of the tetrasaccharide Man-β(1-4)-GlcNAc-β(1-4)-[α(1-6)-Fuc]-GlcNAc, named here sugar D for short, see **Figure 1**, is quite complex and shows a rare event where the ring of the reducing GlcNAc undergoes a change of pucker from the stable ^4^C_1_ chair conformation, to a less stable ^1^C_4_ chair. Similar conformational changes have been reported recently in the literature for another tetrasaccharide, namely Lewis X^39-40^, where the linkage of the GlcNAc to the fucose is a less flexible β(1-3). Although we were able to observe the first ^4^C_1_ to ^1^C_4_ chair transition within the first 100 ns of the simulation, we extended the trajectory to 3 µs to reach convergence. During the 3 µs trajectory the ^1^C_4_ chair conformation is visited 5 times for intervals ranging from 30 ns up to 200 ns. Nevertheless, as shown in **Figure 5,** during the trajectory sugar D is found in the stable ^4^C_1_ chair for over 87% of the time, which in terms of relative populations, the free energy for the ^4^C_1_ to ^1^C_4_ chair transition corresponds to 4.7 kJ/mol at 300 K.

**Figure 5.**
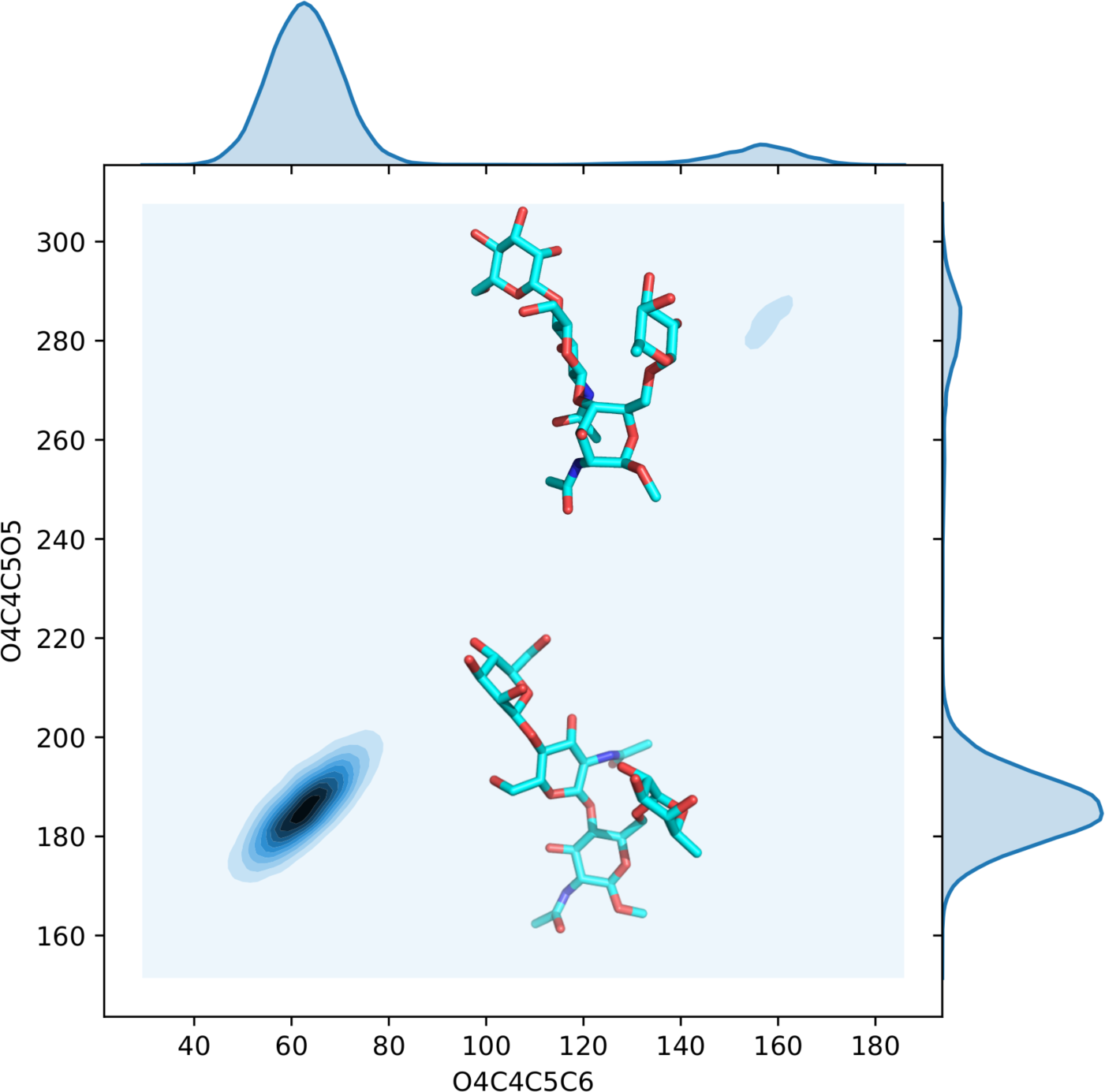
Contour plot showing the conformational propensity of the reducing GlcNAc(1) of sugar D in function of two torsion angles O4C4C5C6 and O4C4C5O5 on the x and y axis, respectively. The structures corresponding to the highest populated ^4^C_1_ chair conformation (87%) and the lowest populated ^1^C_4_ chair conformation (13%) over 3 µs (30,000 data points) are also shown in line with the corresponding histograms on the y axis. Torsion angles are in degrees. 2D contour plots made with seaborn, with sugar D structures rendered with pymol.

## Discussion

The sequence of the N-glycans expressed at Asn 297 has been shown to have a significant effect in modulating the IgG effector function. In particular core-fucosylation, and galactosylation and sialylation of both α(1-6) and α(1-3) arms in biantennary N-glycans have been identified as key players, with different roles in regulating the ADCC function^5, 13, 16^ and the onset of inflammation^22, 24^. In particular levels of galactosylation have been linked to aging^29^ and to the risk of developing conditions such as rheumatoid arthritis^23, 25-27^. The molecular basis underlying the link between the glycan sequence and these different phenotypes is very complex to understand, as it most likely depends on the dynamics and energetics of the interaction between the glycosylated antibody and the different glycosylated FcγRs^19^. To begin to understand these processes from an atomistic level of detail, as a first step we analysed the intrinsic conformational propensity of the biantennary N-glycans most commonly expressed at Asn 297 in the IgG Fc^5, 10^ to determine if and how the monosaccharide sequence affects the N-glycan dynamics in the absence of the protein. Although we do expect that contacts with residues on the IgG Fc surface are likely to affect the conformational equilibrium relative to the isolated glycan, it is has been shown that both α(1-6) and α(1-3) arms retain high degrees of flexibility^28^, thus their intrinsic conformational propensity may play a role in their contact with the receptors. The results of our study based on extensive conformational sampling in excess of 62 µs by MD simulations, has shown that while core-fucosylation and sialylation do not affect the conformational dynamics of the N-glycan as a whole, the effect of galactosylation of the α(1-6) arm is remarkable, as it clearly regulates its conformation. Indeed, we have found that regardless of the sequence of the α(1-3) arm, of core-fucosylation, or of sialylation of the α(1-6) arm, the presence of galactose β(1-4) linked to GlcNAc shifts the conformational propensity of the whole α(1-6) arm from an ‘outstretched’ conformation, shown in **Figure 4** on the left, to a ‘folded-over’ conformation where the α(1-6) arm is stacked against the chitobiose core. The shift in the conformational equilibrium we observe for galactosylated α(1-6) arms is likely due to a more effective interaction in terms of both hydrogen bonding and of stacking with the chitobiose core, regardless of the presence of the core-fucose. This is surprising as fucose does contribute to the hydrogen bonding network by bridging the GlcNAc(2) of the chitobiose to the GlcNAc(7) of the folded-over α(1-6) arm, as shown in Figure 2. Conversely, the dynamics of the α(1-3) arm is more restricted due to the intrinsic rigidity of the Man-α(1-3)-Man relative to the Man-α(1-6)-Man. Nevertheless it undergoes excursions between the two rotamers shown in **Figure 3**. The conformational dynamics of both armss is locally enhanced by the sialic acid due to the intrinsic flexibility of the α(2-6) linkage.

Although the existence of ‘folded-over’ (and variation there-of) and of ‘outstretched’ conformations of the α(1-6) arm have been identified and discussed previously^41-42^, our systematic study highlights the unique dependence of this conformational propensity on the galactosylation of the α(1-6) arm. This behaviour can explain the recent evidence of differential recognition of positional isomers in glycan array screening^43^. As an example, the positional isomers sugar J and L have a galactose on either the α(1-3) or the α(1-6) arm, respectively, which confers a different conformational propensity of the α(1-6) arm. Indeed, in sugar J, shown in **Figure 6a)**, the ‘folded-over’ and the ‘outstretched’ conformations are equally populated during the MD simulations, i.e. 49% and 50%, respectively (note: the 1% corresponds to a ψ value of −99.3°), while in sugar L, shown in **Figure 6 b)**, the ‘folded-over’ α(1-6) arm is the dominant conformation with a relative population of 81% over the simulation time. As a further implication of this behaviour, a different structural propensity shifted towards a ‘folded-over’ α(1-6) arm and an outstretched α(1-3) arm is in agreement with the higher affinity of the sialyltransferase for the α(1-3) galactose, more accessible, in isolated glycans^28^. Interestingly, this selectivity is only moderately changed when the N-glycans are linked to the IgG Fc^28^.

**Figure 6.**
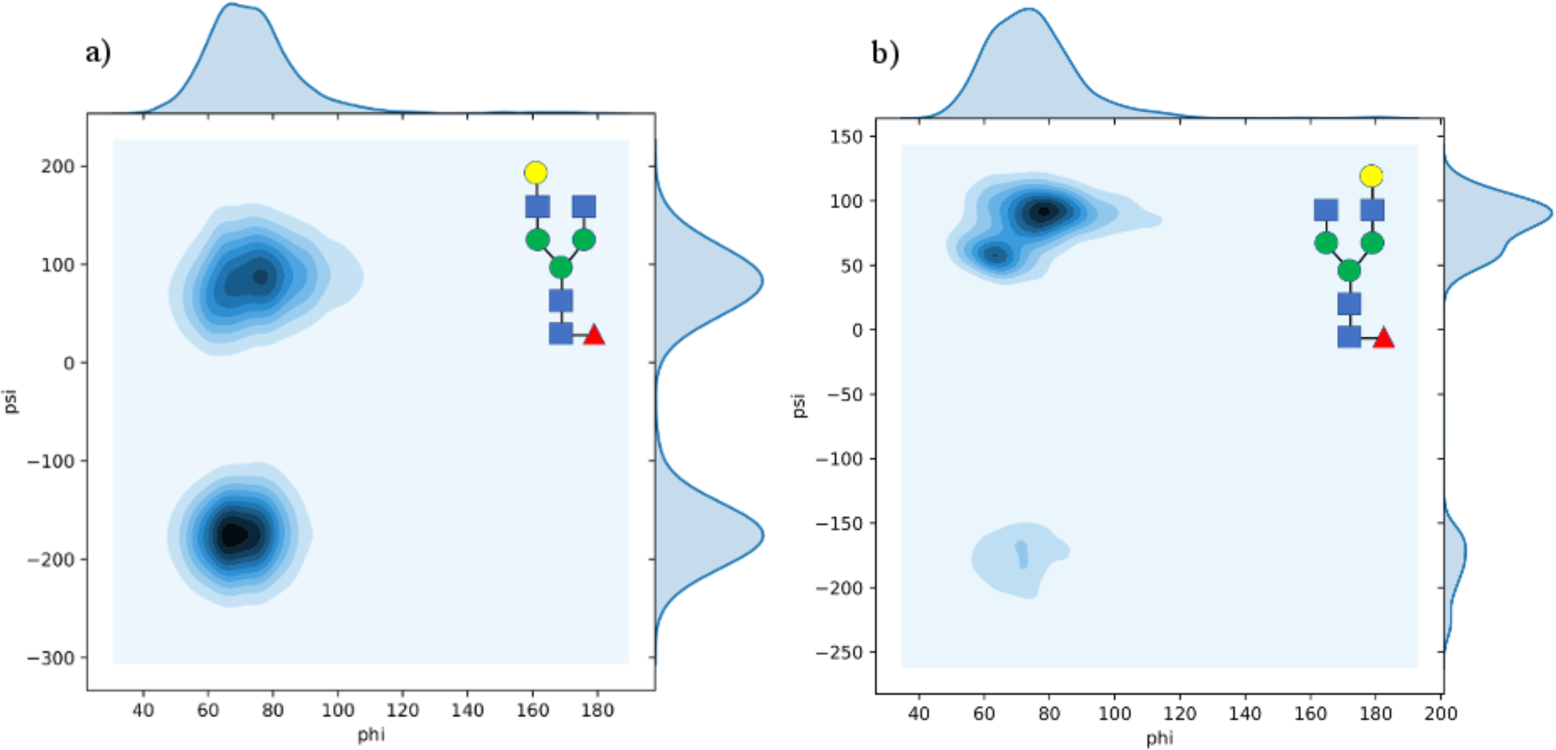
Contour plot showing the different conformational propensities of the α(1-6) arm in sugar J (panel a) and L (panel b) in function of galactosylation. The ‘folded over’ conformation corresponds to a ψ value around 80°, while the outstretched conformation corresponds to a ψ value around −180°. Torsion angles are in degrees. 2D contour plots made with seaborn with 2,500 data points.

## Conclusions

Here we have shown the results of extensive sampling by unbiased MD simulations of the conformational space accessible to increasingly larger biantennary complex N-glycans commonly expressed at the Asn 297 in the IgG Fc region. Our data indicate that while core-fucosylation and sialylation do not affect the overall conformation of the isolated N-glycan as a whole, but contribute to its local dynamics, galactosylation of the α(1-6) arm shifts its conformational equilibrium towards a structure where the arm is ‘folded over’ the chitobiose core. This effect is determined by more effective hydrogen bonding and stacking interactions of the chitobiose with the ‘longer’ galactosylated α(1-6) arm and it is independent of the sequence of the α(1-3) arm, of core-fucosylation and of sialylation of the α(1-6) arm. These results can explain the differential recognition of positional isomers^43^ and with the preference of sialyltransferases for the α(1-3) arm in both, isolated and Fc-linked N-glycans. Currently we are in the process of determining how the dynamics and the conformational equilibria we have discussed here for the isolated biantennary N-glycans are affected by the presence of the IgG Fc protein surface.

## Acknowledgements

The authors gratefully acknowledge the Irish Centre for High-End Computing (ICHEC) for the generous allocation of computational resources. EF would like to thank Dr Frank Fieschi (IBS) for insightful discussions. The John and Pat Hume Doctoral Awards programme at Maynooth University is also gratefully acknowledged for funding.

